# The Human Brain encodes a Chronicle of Visual Events at each Instant of Time

**DOI:** 10.1101/846576

**Authors:** J-R. King, V. Wyart

## Abstract

The human brain continuously processes streams of visual input. Yet, a single image typically triggers neural responses that extend beyond one second. To understand how such a slow computation copes with real-time processing, we recorded subjects’ electrical brain activity, while they watched ~5,000 rapidly-changing images. First, we show that each image can be decoded from brain activity for ~1 sec, and demonstrate that the brain simultaneously represents multiple images at each time instant. Second, dynamical system modeling reveals that these sustained representations can be explained by a specific chain of neural circuits, which consist of (i) a hidden maintenance mechanism, and (ii) an observable update mechanism. Third, this neural architecture is localized along the expected visual pathways. Finally, we show that the propagation of low-level representations across the visual hierarchy is a principle shared with deep convolutional networks. Together, these findings provide a general neural mechanism to simultaneously represent successive visual events.

**Significance:** Our retina are continuously bombarded with a rich flux of visual input. How our brain continuously processes such visual stream is a major challenge to neuroscience. Here, we developed techniques to decode and track, from human brain activity, multiple images flashed in rapid succession. Our results show that the brain *simultaneously* represents multiple *successive* images at each time instant. A hierarchy of neural assemblies which continuously propagate multiple visual contents explains our findings. Overall, this study sheds new light on the biological basis of our visual experience.

## Introduction

The human visual system is continuously bombarded with a flux of visual input. To interact with its environment, our brain must, therefore, transform at each time instant these visual events into abstract representations (Tenenbaum et al., 2011), track their relative motions (Born and Bradley, 2005; Bruce Goldstein and Brockmole, 2016) and resolve countless ambiguities (Knill and Pouget, 2004).

However, to decipher the visual system, electrophysiology and neuroimaging studies have primarily focused on the brain responses to static images (although see (Marti and Dehaene, 2017; Nishimoto et al., 2011; VanRullen and Macdonald, 2012; van Vugt et al., 2018)). The resulting studies consistently show that flashing an image onto the retina leads to cortical responses that can last up to one second (Carlson et al., 2013; Cichy et al., 2014; King et al., 2016). The timing and location of these neural activations are consistent with a hierarchical inference network (Cichy et al., 2014; Friston, 2010; Poggio and Anselmi, 2016; Yamins et al., 2014) and suggest that the primary visual cortex encodes low-level representations (e.g. luminosity and orientations of visual edges), while the infero-temporal and dorso-parietal cortices encode abstract representations (e.g. the presence of a face in the visual field).

However, these single-stimulus studies highlight a fundamental paradox: the duration of visual processing (~1,000 ms) can largely exceed the duration of sensory stimulation (e.g. 17 ms (King et al., 2016)). Consequently, it is unclear how, at the macroscopic level, the brain copes with streams of sensory inputs. Are past visual contents simply replaced by incoming stimulations or can multiple snapshots of the past be simultaneously represented in the brain activity? Are visual areas continuously feeding their representations up the visual hierarchy, or do they only transmit the representations that need to be updated? Are low-level representations maintained in early sensory areas and if so, how do associative cortices bind and integrate these continuously changing features?

To address these questions, we recorded the electrical brain activity of human subjects with electroencephalography (EEG), while they watched ~5,000 parametrically-controlled images flashed in rapid succession. First, we show with decoding analyses that the human brain *simultaneously* represents multiple images that have been presented *sequentially*. Second, we formalize with a dynamical system framework, the neural architectures that can in principle maintain successive images. Third, we show with temporal generalization and source analyses that each image triggers an update signal that propagates across the visual hierarchy. Finally, we show that these feedforward signals accumulate increasingly abstract features across the visual hierarchy.

## Results

### Successive visual stimuli can be simultaneously decoded at each time instant

Capitalizing on both neuroscientific studies (Stocker and Simoncelli, 2006) and computer vision practices (Wang et al., 2016), we hypothesized that the brain responses to visual streams may represent the instantaneous visual content (i.e. the image presented at a given time point) and/or the instantaneous optical flow (the change between two successive images, Fig. 1A). By definition, these two features cannot be easily disentangled in single-stimulus studies. We thus recorded the brain electric activity of fifteen healthy subjects with EEG, while they watched rapid sequences of randomly-oriented Gabor patches, grouped into 8-stimulus trials (Fig. 1. B-C).

**Figure 1.**
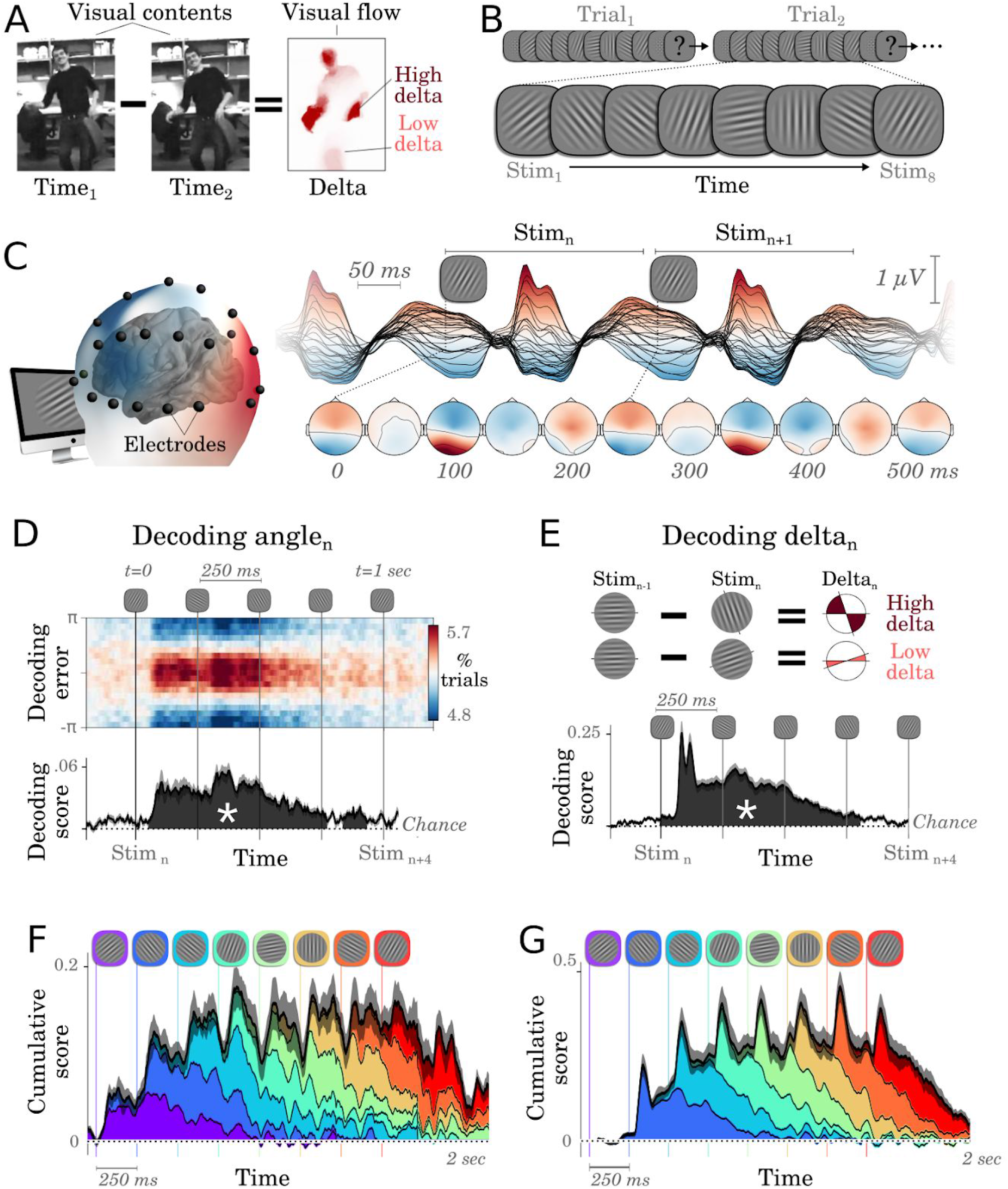
Successive images are simultaneously represented in brain activity. **A.** Visual contents refers to what is in the image at time *t*. Visual flow refers to the amount of change between t_1_ and t_2_. **B.** Subjects watched ~5,000 randomly oriented Gabor patches, flashed every 250 ms and grouped into 8-item sequences separated by masks. Each sequence ended in a two-alternative forced choice where subjects indicated whether stimuli fell, on average, closer to the cardinal or diagonal axes. **C.** Brain activity was recorded with EEG. Each line shows the average response evoked by the stimuli **D.** Top. Distribution of single-trial decoding error of Stim_n_ as a function of time relative to the onset of stimulus n. *Bottom*. Time-course of the corresponding decoding score. The shaded regions (with an asterisk) indicate significant decoding across subjects (cluster corrected). **E.** Decoding scores of visual flows (approximated as the absolute angular difference between successive stimuli) as a function of time relative to stimulus onset. **F.** Cumulative decoding scores (black) and the contribution of each of the eight successive stimuli (color-coded by position in the 8-item sequence), as a function of time relative to the sequence onset (chance=0). **G.** Similar to F for cumulative delta decoding scores. In panels C-G, the vertical lines mark the onsets of each stimulus. Error bars indicate the standard error of the mean across subjects.

We then tested whether these EEG signals linearly correlated with (i) the orientation of each stimulus *n* (angle_n_), as well as with (ii) the absolute change between successive stimuli (delta_n_ = |angles_n+1_ - angle_n_|; deltas and angles are orthogonal by design). To this aim, multivariate linear regressions (*W*_*t*_) were fit at each time sample *t* relative to the onset of each stimulus in order to predict the sine and cosine of its angle given the voltage of all EEG electrodes. We then assessed with cross-validation whether these predicted angles correlated with the true stimulus angles. This “angle decoding” was significantly above chance across subjects between ~50 and ~950 ms after stimulus onset (cluster-corrected effects across subjects illustrated in Fig. 1D). These results demonstrate that the long-lasting brain responses typically reported in single-stimulus protocols can also be observed when an image is presented in a rapidly changing visual stream.

We applied an analogous analysis to decode the change of orientations between successive pairs of stimuli (delta) in order to assess whether and when brain activity represented optical flows. The corresponding “delta decoding” time-course was similar to the angle decoding time course (Fig. 1E). However, encoding analyses revealed that the variance of EEG evoked responses was much better accounted for by stimulus deltas (R=0.20) than by stimulus angles (R=0.05, delta-angle: p<0.0001). In line with recent practices in artificial vision, these findings suggest that the brain not only represents the content of individual images, but also represents the optical flows between images.

Overall, beyond *what* is represented in the brain, we were surprised by *how long* these representations lasted. In particular, decoding analyses independently assessed for each successive stimulus position within a visual stream revealed that multiple stimuli are simultaneously represented at each time sample (Fig. 1. F-G). On average, between two and five angles and between two and four deltas can be simultaneously decoded at each time point, although with a rapid loss of precision as time passes (linear regression between scores and stimulus distance: β_Angles_=0.007, p=0.001; β_Delta_=0.028, p<.001). To verify that these sustained representations are not trivially induced by subjects’ task (i.e. reporting whether, on average, stimuli fell closer to the cardinal or diagonal axes), we show that (i) the decoding of the task variable specifically rises at the end of each 8-item sequences in the beta frequency band, and that (ii) the maintenance of low-level representations is not necessary to perform subjects’ task (Fig. S4).

### Distinct neural architectures can maintain successive stimuli

How does the brain continuously maintain such a long a succession of visual stimuli? A variety of cellular and/or population mechanisms can be proposed. For example, neuronal adaptation induced by the slow return of sodium and potassium currents that follow a spike, temporarily diminishes the expected spiking activity (Fig. 2A), which would thus result in a re-activation of decoding performance after stimulus offset (Fig. 2B). Alternatively, recurrent connections between excitatory and inhibitory neurons can lead to similar dynamics at the population level. In both case the corresponding dynamical system consists of (1) an observable unit *x* (e.g. membrane potential or mean-field activity of post-synaptic potentials (psp) to excitatory neurons) and (2) a hidden unit *y* (e.g. adaptive ion currents or psp to inhibitory neurons) connected in a negative feedback loop. More generally, various types of feedback loops (Fig. 2B) and feedforward propagations (Fig. 2.C-D) can be implemented to maintain sensory representations in the EEG activity, even after the offset of the stimulus.

**Figure 2.**
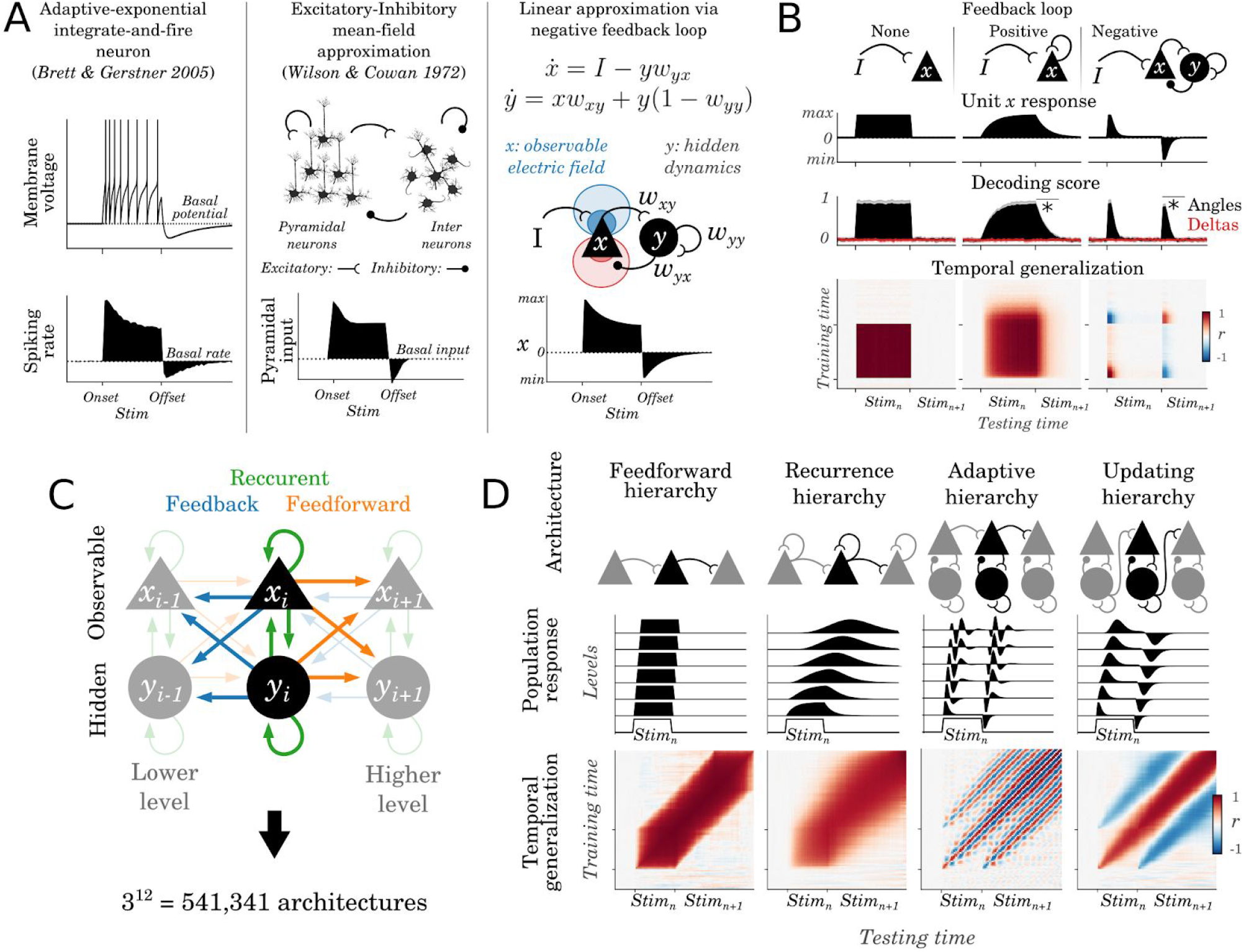
Dynamical system framework. **A. Left.** The membrane potential (top) and the expected spike rate (bottom) of an AdexpIAF neuron in response to a sustained input (onset and offset indicated by the ticks); the dotted lines indicates the basal activity at rest. **Middle.** In an excitatory/inhibitory population of neurons stimulated with an input, the alignment of pyramidal dendrites leads their post-synaptic potentials (psp) to be detectable from distant electrodes, whereas inter-neurons’ psp are not detectable with EEG. **Right.** A linear dynamical system, composed of two units (*x*: observable and *y*: hidden) connected in a feedback loop can approximate adaptive neurons or E/I balance responses: *x* captures an observable variable (e.g. the electric field associated with spiking activity or pyramidal psp) whereas *y* captures a hidden variable (e.g. ion-currents associated with adaptation or inhibitory psp). **B.** Columns illustrate the predictions of positive, negative and no feedback loop circuits respectively. The top black line illustrate the activity of an observable unit (x) in response to a stimulus (onset and offset marked by ticks). Decoding scores of stimulus angles (black) and deltas (red) from the simulated population x tuned to stimulus angles. The asterisks highlight whether Stimn can be decoded after its offset. The temporal generalization (TG) matrices correspond to the decoding scores of each decoder trained at time t and tested at all time samples. **C.** More complex networks can be generated by hierarchically connecting feedback loops with one another. Arrows indicate connections within or between the levels of such hierarchy. **D.** Examples of plausible hierarchies, together with (top) the dynamics of their observable units (*x*) at each hierarchical levels (black lines) in response to a brief stimulus and (bottom) the corresponding temporal generalization matrices.

Temporal generalization (TG) analyses can be used to distinguish such dynamical systems. TG consists in quantifying the extent to which a linear decoder trained at time *t* (across all EEG channels) can accurately decode representations at time *t’* (King and Dehaene, 2014; Stokes et al., 2013). For example, a positive feedback loop input with a brief stimulus leads to a ‘square’ TG matrix that extends beyond stimulus offset (Fig. 2C). By contrast, a negative feedback loop input with the same stimulus leads to a rapid decrease of decodable activity (potentially down to chance level) followed by below-chance decoding after stimulus offset. TG can also be used to differentiate more complex dynamical systems (Fig. 2D). For example, a strictly feedforward architecture leads all linear decoders trained at a given time sample to generalize for a constant (but temporally shifted) time period. By contrast, a chain of positive feedback loops (‘Recurrence Hierarchy’) leads linear decoders to generalize over increasingly longer time periods. Depending on the type of connection between units, a chain may lead to oscillatory activity (e.g. ‘Adaptive Hierarchy’, Fig. 2D) or to positive and negative traveling waves triggered by stimulus onsets and offsets respectively (Fig. 2D ‘Updating hierarchy’).

### Visual representations propagate across a neural network

Do the hand-picked dynamical systems illustrated in Fig. 2 match the spatio-temporal characteristics of the brain responses to visual streams? To address this issue, we separately implemented TG for each subject and verified with non-parametric cluster-level testing across subjects whether each angle and delta decoder trained at a given time sample was able to generalize over all time samples (Fig. 3 A-B). We then tested whether the significant TG clusters matched the predictions of the models illustrated above. The analyses revealed four main findings. First, both angle and delta TG matrices were characterized by a ~900 ms long (above-chance) cluster around the TG diagonal (angle: p<0.001; delta: p<0.001). This diagonal pattern invalidates the predictions of non-hierarchical circuits: i.e. positive or negative feedbacks alone do not lead to diagonal TG. Second, the angle and delta decoders successfully generalized, on average, over 75 ms (s.d.=68 ms) and 129 ms (s.d.=54 ms) respectively. These durations are shorter than both (i) the time period during which angles and deltas are decodable (~900 ms, shaded areas in Fig. 1 D-E) and (ii) the stimulus duration (233 ms). These brief generalizations, therefore, invalidate the predictions of the “feedforward” and “positive feedback loop” chains, as both architectures predict decoders to generalize for a time period longer than the stimulus presentation. Third, these generalization periods consistently increased in duration as a function of training time (angles: r=0.44, p<0.001; deltas: r=0.47, p=0.003, Fig. 3A) as predicted by the chains of feedback loops. Fourth, the central positive decoding pattern was surrounded by two below-chance diagonals (p<0.01, for both angles and deltas) time-locked to the stimulus offset (blue areas in Fig. 3 A-B) as predicted by the chain of negative feedback loops. Supplementary analyses show that the reversal of neural responses after stimulus offset partially account for the long-lasting decoding angle and delta scores (Fig. S1).

**Figure 3.**
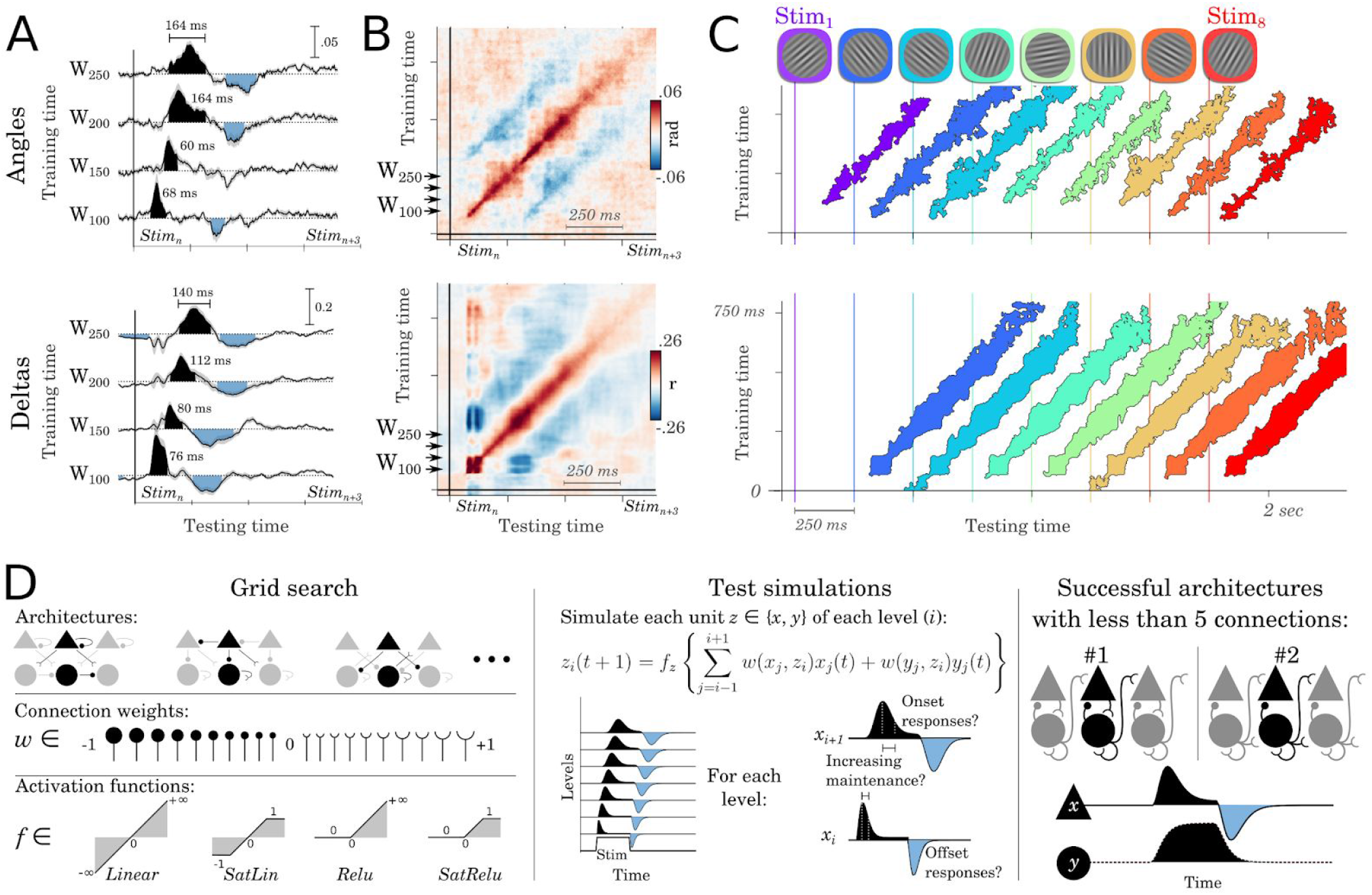
The spatio-temporal dynamics of representations reveal an ‘Updating Hierarchy’. **A.** Examples of TG for angle (top) and delta (bottom) decoders trained at 100, 150, 200 and 250 ms after stimulus onset, and tested across all time samples. The shaded areas indicate significant generalization, cluster-corrected across subjects. Time annotations indicate the duration during which each decoder significantly generalized. **B**. Full TG matrices for angle (top) and delta (bottom) decoders. Blue areas indicate below-chance generalizations. **C.** TG scores for each of the eight successive stimuli. Colored areas indicate the above-chance generalizations, cluster-corrected across subjects. **D. Left**. Grid-search analyses across architectures, connections weights (*w*) and activation functions (*f*) led to simulate >1.5 billion hierarchical models. **Middle**. Each of them was tested on its ability to generate dynamics qualitatively similar to those obtained empirically: i.e. characterized by onset and offset responses whose durations increased across levels. **Right**. Two architectures captured these dynamics with no more than four connections. The plain line illustrates a representative example of an observable unit (x). The dotted line illustrates a representative example of the hidden unit (*y*).

To test whether these TG patterns were consistently observed for each stimulus, we independently decoded the angle and delta as a function of the position of the stimuli within the 8-item sequence. The corresponding TG analyses resulted in a series of parallel and non-overlapping diagonals (Fig. 3C). Overall, these results suggest that successive visual events are simultaneously encoded in brain activity thanks to a rapid propagation of their corresponding representations along a long chain of negative feedback loops.

The models presented in Fig. 2 illustrate a small set of possible neural architectures, as defined by the signs of the connections between units (i.e. excitatory, none, or inhibitory, Fig. 2D). To systematically investigate which architecture could account for subjects’ brain activity, we performed a grid search analysis by (i) systematically varying each connection (min=−1, max=1, step=0.05) of each architecture as well as by (ii) testing four monotonic activation functions (Linear, ReLu, SatLin, SatRelu) for each unit type (*x*: observable and *y*: hidden). Each of the tested networks consisted of a 10-layer hierarchy of *x* and *y* units inter-connected with up to four recurrent, four feedforward and/or four feedback connections (Fig. 2C). The resulting models were evaluated on their ability to generate dynamics qualitatively similar to those obtained empirically (see Supplementary methods: Model search). Approximately 1.5 billion models were tested and spanned the 531,441 distinct architectures. The model search showed that only two architectures presented dynamics qualitatively similar to subjects’ EEG with no more than four connections. The first architecture corresponds to the ‘Updating hierarchy’ (Fig. 2D). The second architecture is also a hierarchy of negative feedback loops, within which *x* units are epiphenomenal. Importantly, both architectures predict that the hidden units *y* maintain stimulus information over time, whereas the observable units *x* mark the update of these representations.

Overall, these results suggest that angles and deltas representations are paradoxically maintained by a biological mechanism undetectable with macroscopic electric brain activity. The long-lasting decoding scores of these two types of representations appear to result from an “update” signal that propagates across a neuronal chain recruited after the onset and the offset of the stimuli. We discuss in the supplementary materials how this dynamical system may partially relate to adaptation, and highlight potential differences (Fig. S1-S2).

### The propagation of representations is localized along the ventral and dorsal visual pathways

Are these traveling waves confined to early visual cortices? To identify the anatomical substrates of the above coding dynamics, we analyzed the spatial patterns of angle and delta representations with encoding analyses. For both angle and delta analyses, the spatial patterns peaked around occipital electrodes and subsequently propagated to anterior electrodes (Fig. 4. A, D). The minimum norm estimates of these scalp topographies suggest that earliest neural responses coding for both angles and deltas are generated in the early visual cortex, and are subsequently followed by responses in the infero-temporal and dorso-parietal cortices (significant clusters summarized in Fig. S3). Furthermore, neural responses coding for deltas reached significance in the superior frontal cortex around 600 ms after stimulus onset. Finally, a full decomposition of the brain responses into their relative peak latencies and peak amplitudes confirmed the overall occipito-frontal propagation of the representations of stimulus angles (r=0.15, p<0.01) and stimulus deltas (r=0.30, p<0.001, Fig. 4.B-C, E-F). Overall, although EEG source reconstruction should be interpreted with caution, our estimates extend the classic notion that stimuli trigger a macroscopic traveling wave across the visual pathways (Riesenhuber and Poggio, 1999; van Vugt et al., 2018) by showing that this traveling wave carries low-level visual information.

**Figure 4.**
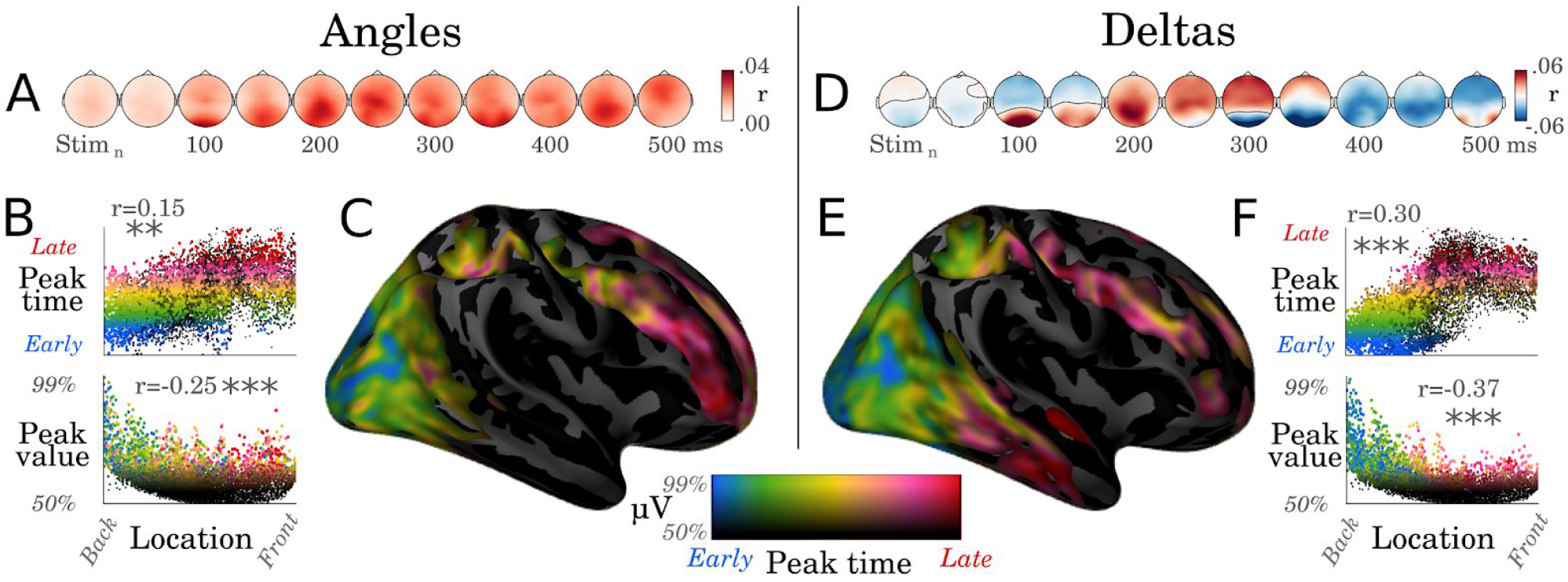
Visual representations propagate from sensory to associative cortices. **A.** Correlation scores resulting from encoding analyses, trained to predict the EEG activity from the sine and cosine of the stimulus angles. **B.** Each dot corresponds to a source estimated from the EEG coding topographies with a minimum norm estimation. The x-axis corresponds to the source location along the postero-anterior direction. The y-axis either indicates the relative timing of the peak activity in each source (Top panel) or the intensity of this peak (Bottom panel). Asterisks indicate statistical significance (**: p<.01, ***: p < 0.001) **C.** Same data as B but plotted on the cortical surface. Colors indicate both the peak amplitude (e.g. black: amplitude = median amplitude across sources) and the peak latency (e.g. blue: peak within the 5% percentile of the earliest responses across sources, red: peak beyond 95% percentile). **D.** Correlation coefficients between deltas and EEG amplitude. **E- F.** Analogous analyses tp B-C applied to the brain responses coding for the changes between successive stimuli (Delta).

### Low-level representations are accumulated across the visual hierarchy

The above results suggest that low-level visual representations, such as the orientation of a flashed Gabor patch, are not confined within the primary visual cortices, but are in fact encoded at multiple levels of the visual hierarchy. This finding originally appeared to us at odds with the notion that the visual hierarchy represents increasingly-complex features (Fig. 5. A-B). We thus hypothesized *ad hoc* that low-level features may be “accumulated” across the visual hierarchy, such that the highest levels of the hierarchy still represent low-level features (Fig. 5B). To test this hypothesis, we first generated increasingly-complex features for each stimulus by repeatedly applying a normalized rectification 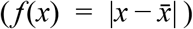 over the stimulus feature (i.e. low-level feature = *f*(angles); middle-level=*f*(*f*(angles)); high=*f*(*f*(*f*(angles))); Fig. 5C). Second, we verified that these increasingly-complex features were decoded in the expected order across the layers of VGG-19 (Simonyan and Zisserman, 2014), a deep convolutional neural network optimized for natural image categorization which has recently been shown to generate representations similar to those of the visual system (Hong et al., 2016). The results confirmed that low-level features are linearly decodable from the first layer, whereas high-level features only become decodable from deeper layers (Fig. 5D). Critically, and contrary to our original intuitions, low-level features remained decodable down to the deepest layers. In other words, a computational hierarchy optimized for visual processing may plausibly benefit from partially propagating low-level representations up to its highest computational stages.

**Figure 5.**
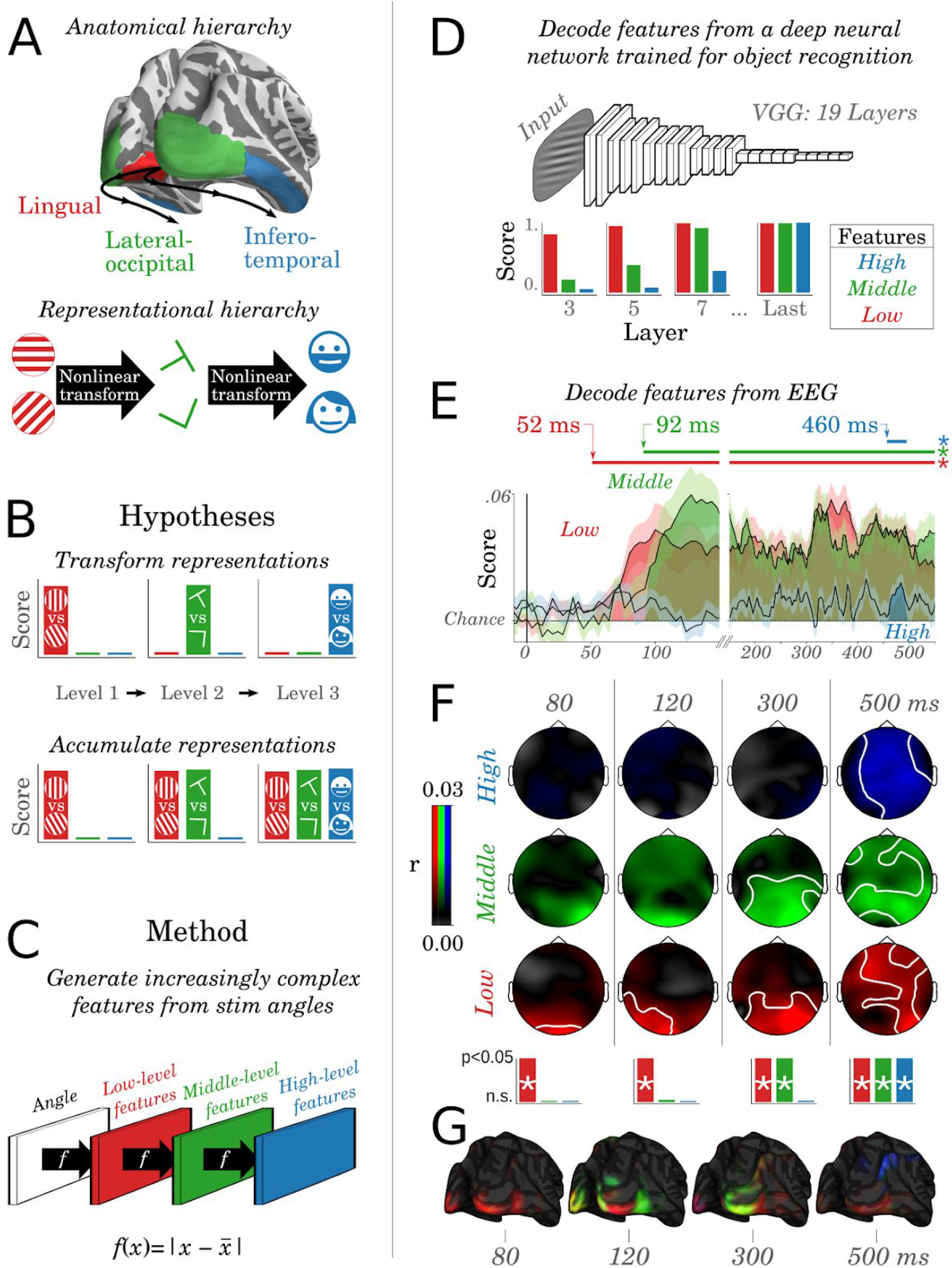
Testing the ad-hoc hypothesis of an accumulation of low-level features across the visual hierarchy. **A.** Primary visual cortices (red) are thought to generate low-level features (e.g. orientations of gabor patches), while higher visual areas (green) and associative cortices (blue) generate increasingly complex features (e.g. from the presence of specific visual edges to the presence of specific faces) through nonlinear activation functions **B.** Hypotheses’ predictions. The histograms illustrate the expected decodability of low-, intermediate- and high-level features across the hierarchy **C.** Increasingly complex features are generated from the stimulus angles by repeatedly applying a nonlinear rectifying function *f*. **D.** Low, middle and high-level features were decoded from the activations of each layer of VGG-19, a deep feedforward neural network trained to identify objects from natural images. **E.** Decoding scores (r) as a function of time for the low- (red), middle-(green) and high-level features (blue). Horizontal lines mark significant cluster-corrected decoding scores. **F.** Encoding correlation scores of the EEG spatio-temporal topographies across features (rows) and time (columns). White lines indicate significant spatio-temporal clusters across subjects (all clusters p<0.001). The asterisks indicate the presence of a significant spatial cluster at these time samples. **G.** The brain source estimates of F.

To test directly test whether this accumulation of features could be observed in brain activity, we applied these “increasingly-complex” analyses to our electrophysiological recordings. The results revealed that visual features started to be decodable in the expected order: low-level feature reached significance at 52 ms (p<0.001), the middle-level cluster started at 92 ms (p<0.001) and the high-level cluster started at 460 ms (p=0.028). Overall, the decoding of the high-level feature was very low (all R<.02, Fig. 5E), but was associated with a consistent (p<0.001) spatio-temporal cluster of electrodes across subjects between 300 and 550 ms after stimulus onset (Fig. 5F). The anatomical sources estimated from these topographical patterns followed the anatomical organization reported in Fig. 4: i.e. they first peaked in the primary visual cortex, expanded to the lateral occipital areas and eventually reached the associative cortices (Fig. 5G). Together, these results suggest that low-level features propagate and accumulate across the visual hierarchy, and thus confirm that multiple representations, characterizing distinct moments of time, can be simultaneously encoded in the overall brain activity.

## Discussion

Three main results reveal how the human brain continuously processes its visual influx. First, we show that the ~1 second-long representations typically observed in single-stimulus studies (Carlson et al., 2013; Cichy et al., 2014; King et al., 2016; VanRullen and Macdonald, 2012) can also be observed in the context of visual streams. This paradoxical discrepancy between the timing of sensation and the timing of neural processing shows that the visual system can continuously buffer and update multiple snapshots of the past. In addition, these brain responses not only represent the *content* of visual input but also, and in fact predominantly represent the *change* between successive images. Overall, these results echo recent practices in artificial vision. Indeed, deep neural networks trained to process videos are typically based on two parallel pathways dedicated to optical content and to optical flow respectively and are input with multiple successive snapshots extracted from videos (Wang et al., 2016). Our results thus suggest that the similarities between artificial and biological neural networks observed processing still images (Güçlü and van Gerven, 2015; Hong et al., 2016; Kriegeskorte, 2015) may extend to those processing videos.

Second, both visual content and visual flow appear to simultaneously propagate across a long chain of neuronal populations, source-localized along the expected visual pathways. This second set of results extends the findings of single-stimulus studies (Cichy et al., 2014; Hung et al., 2005; Lamme and Roelfsema, 2000) to visual streams. Importantly, dynamical system modeling and temporal generalization suggest that the traveling waves elicited at stimulus onset and offset reflect a series of transient signals triggered by the change between images. This feature, partially accounted for by sensory adaptation (Fig. S2) is reminiscent of predictive coding architectures (Friston, 2018; Rao and Ballard, 1999). Our model search further show that visual representations are maintained, within each level of a macroscopic hierarchy, thanks to biological mechanisms undetectable with EEG. Overall, the presently identified dynamical system suggests that dynamic coding (Stokes, 2015) and activity-silent maintenance (Wolff et al., 2017) may reflect the update and the maintenance mechanisms of a unique neural architecture.

Third, our results suggest that low-level visual representations remain encoded in multiple levels of the visual hierarchy. This phenomenon extends similar findings recently observed in the macaque’s infero-temporal cortex in response to static images (Hong et al., 2016) to the processing of visual streams in humans. Furthermore, such propagation of low-level features reminded us of the success of Residual networks, which outperform deep convolutional networks by explicitly accumulating features across layers (He et al., 2016). Here, we show that this accumulation may, in fact, already be approximated by trained deep convolutional ANNs.

Finally, our EEG study faces several limitations. First, it is based on artificial visual streams with a fixed presentation rate. It thus only provides a lower-bound estimate of the number of successive images simultaneously encoded in brain activity. In particular, it is unclear whether the relevance of these images with regard to the task lead subjects to maintain each of them for a long time-period.

Overall, the macroscopic dynamical system revealed in this study implies an important computational consequence. Specifically, the redundant encoding of visual features across the hierarchy allows the prefrontal and parietal cortices, which receive inputs from the whole visual hierarchy (Dehaene and Changeux, 2011), to instantaneously bind both (i) multiple levels of representations and (ii) multiple successive images with a linear readout. We thus look forward to test this hypothesis with single cell recordings of the associative cortices. More generally, characterizing the anatomical, dynamical and representational properties of the visual system paves the way to understand how single-cells (van Vugt et al., 2018) and cortical microcircuits (Gavornik and Bear, 2014) are collectively organized to continuously transform the avalanche of sensory input into a coherent stream of mental representations.

## Materials and Methods

### Subjects

Sixteen healthy adults (age range: 18-25 years), all with normal or corrected-to-normal vision and no reported history of neurologic or psychiatric disorders, were recruited from the University of Oxford. Subjects provided written consent before participating in the experiment and received £30 in compensation for their participation. Approximately £5 of bonuses could be additionally obtained depending on subjects’ performance on the two-alternative forced-choice task. The study followed local ethics guidelines. The data from one participant was excluded because of eye artifacts. The investigation of subjects’ decisions at the end of each sequence is reported in (Wyart et al., 2012).

### Procedure

The stimuli were displayed on a CRT monitor (1024 × 768 pixels, refreshed at 60 Hz) placed approximately 80 cm away from subjects’ eyes, and controlled with the MATLAB (The Mathworks, Natick, MA) Psychophysics-3 toolbox (Brainard, 1997; Pelli, 1997). Each trial consisted of a sequence of eleven successive visual stimuli flashed every 250 ms. The first two items and the last item were task-irrelevant masks, generated from the average of four cardinal and diagonal Gabor patterns and were not considered in the present analyses. The purpose of these masks was to increase the homogeneity across the stimuli: e.g. the first Gabor patch would already be presented in a stream. The remaining eight stimuli (hereafter referred to as the 8-item sequences) were Gabor patches with fixed contrast (50%), diameter (4 degrees of visual angle), spatial frequency (2 cycles per degree of visual angle), and envelope (Gaussian with a s.d. of 1 degree of visual angle). Stimulus orientations, however, varied following a uniform distribution across all trials. One blank frame (16.7 ms) was introduced before the onset of each stimulus to avoid visual ‘tearing’ artifacts. The inter-trial interval was 1.250 s (±250 ms). The experiment consisted of 672 trials, divided into seven sessions of 96 trials, and thus consisted of a maximum of 5,376 usable brain responses to oriented stimuli over approximately one hour of recording.

### Task

To ensure subjects paid attention to the visual stimuli, they were asked to report after each 8-stimulus sequence whether the orientation of the stimuli was, on average, closer to (i) the cardinal axes (horizontal or vertical) to (ii) the diagonal axes (45° or 135°). Responses were given by pressing their left or right index finger on either of the right or left CTRL buttons (e.g., diagonal = left finger and cardinal = right finger), with a response-mapping counterbalanced across subjects. This two-alternative-forced choice task was adapted for each subject in order to homogenize attention and performance across subjects. Specifically, the orientation of each Gabor patch was distributed uniformly across all trials but drawn, within each sequence, from a probability density function adapted for each subject’s performance. To this end, subjects performed a practice and a titration session before the main experiment in order to estimate their 75% accuracy psychophysical threshold with an adaptive staircase procedure (Kaernbach, 1991). This psychophysical threshold served to determine three evenly-spaced difficulty levels. For example, an easy sequence would consist of stimuli whose orientations tend to fall close to the cardinal axes, whereas a hard sequence would consist of stimuli whose orientations tended to fall in between the cardinal and diagonal axes. Easy and difficult trials (1/3 of all trials each) had a categorization sensitivity of 2.12 (SEM: ±0.18) and 1.00 (SEM:±0.09) respectively. Neutral trials (1/3 of all trials) were associated with a pseudorandom feedback, positive on 60% of neutral trials. Additional behavioral and brain correlates of subjects’ decision are reported in (Wyart et al., 2012), and are orthogonal to the visual contents and visual flows presently studied as detailed in the supplementary analyses and figures (Fig. S4).

### EEG acquisition and pre-processing

EEG signals were recorded with a 32-channel Neuroscan system and a Synamps-2 digital amplifier. In addition, the horizontal and vertical electrooculograms (EOG) were recorded with four bipolar-mounted electrodes. Electrode impedances were kept below 50 kΩ. EEG signals were recorded at 1,000 Hz and high-pass filtered online at 0.1 Hz, and later low-pass filtered at 40 Hz, down-sampled to 250 Hz, segmented from 500 ms before the onset of the first stimulus (the pre-mask) to 1 s following the offset of last stimulus (the post-mask). These epochs were visually inspected (i) to remove trials containing non-stereotypical artifacts, and (ii) to identify artifacted electrode(s). In total three participants had a single bad electrode, which was consequently interpolated to the weighted average of neighboring electrodes.

### Analyses

The stimulus orientation is here used to test whether brain activity represents (i.e. linearly correlates with) the visual content present on the retina at a given instant. Because stimulus orientation is circular, the encoding and decoding analyses of stimulus angles are based on the linear regression of (i) the EEG and (ii) the sine and cosine of the stimulus angles (horizontal = 0 rad; vertical = 2π rad). By contrast, the change of orientation (delta=|angle_n_ - angle_n−1_|) is here used as a way to probe whether brain activity codes for visual flow (Fig. 1A). The first Gabor patch of the sequence was ignored from delta analyses.

### Decoding

Multivariate linear decoding models were implemented following a three-step procedure: fitting, predicting, scoring within subjects. Specifically, for each subject separately we fitted an ordinary least square regression across all EEG channels recorded at a given time sample *t* relative to the onset of stimuli *n*, in order to predict the features of stimulus n (i.e. angle & delta) following the methods of (King et al., 2016). Each decoder thus consisted of a linear spatial filter (*W*):

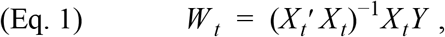

where *X* corresponds to all EEG electrodes recorded at time t after the onset of stimuli and *Y*_*n*_ corresponds either (i) to the delta between stimuli *n* and stimuli *n−1*, or (ii) to the sine and cosine of stimuli n. Each spatial filter W_t_ was then used to predict the angles and the deltas of out-of-sample EEG data (see cross-validation):

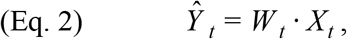

where *Ŷ*_t_ is either (i) the estimated angle or (ii) the estimated delta of each out-of-sample EEG recording X at time t.

Finally, delta decoding scores were summarized with a Pearson correlation coefficient R between the out-of-sample delta predictions and the corresponding true deltas. Angle decoding scores were summarized by computing the angular difference between the true angles (α) and the out-of-sample angular predictions 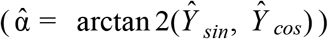. For clarity purposes, this angular error is reported throughout the manuscript as an angular score (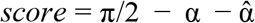; chance = 0). The decoders were trained within each subject with a cross-validation scheme across all stimuli independently of their position in a given sequence. To interpret the decoders, we transformed the spatial filters W into spatial patterns P following the Haufe’s method (Haufe et al., 2014):

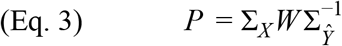
where ∑_X_ and ∑_Ŷ_ refer to the empirical covariances of *X* and *Ŷ* respectively.

### Temporal generalization (TG)

TG analysis consists in testing the ability of each temporal decoder to generalize over all time samples (King and Dehaene, 2014; Stokes et al., 2013). Specifically, each decoder trained at a given time *t* is scored on its ability to decode brain activity from a time *t’*. A significant temporal generalization suggests that the brain activity patterns observed at time *t* are partially similar to those observed at time *t’*. Because our decoders are linear, and because the EEG activity reflect a linear projection of the neuronal activity onto our sensors, a significant generalization suggests that a similar population of neurons is activated at time *t* and *t’*. *Vice versa*, if two decoders trained at *t* and *t’* respectively can both be used to decode the EEG activity at the respective training time, but fail to cross-generalize with one another, then this suggests that the populations of neurons activated at *t* and *t’* significantly differ. Consequently, TG results in a training by testing time matrix which can be used as a method to characterize the spatio-temporal dynamics of the coding response (King and Dehaene, 2014; Stokes et al., 2013).

### Encoding

To assess the extent to which visual content and visual flow comparatively predict subjects’ EEG, we applied encoding analyses for each subject separately. Specifically, we fit, for each electrode and each time-sample separately, an ordinary least-square regression to predict the amplitude of each electrode at each time sample from our experimental variable *Y*:

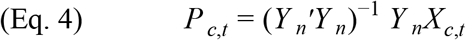

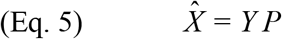

We summarize the encoding results with the Pearson correlation coefficient R obtained for each electrode and each time sample relative to the stimulus onset, between the true voltage and the voltage predicted by our model. Analogous encoding analyses were applied for Fig. 5F. Specifically, we trained, for each subject, at each time sample and for each electrode independently, a linear model to predict the EEG evoked activity given the low, middle or high level features (see Modeling representational structure) separately. Using cross-validation, we assessed whether these predictions correlated with the actual EEG voltage.

### Cross-validation

Decoding and encoding analyses were implemented within an ad-hoc stratified KFold cross-validation procedure split across 8-item sequences. Specifically, cross-validation was designed to ensure that two stimuli from the same 8-item sequences never appeared both in the training and testing sets.

### Source estimates

The locations of the neural sources corresponding to effects observed at sensor levels were estimated following MNE source reconstruction pipeline (Gramfort et al., 2014). The noise covariance was estimated from the 200 ms baseline activity preceding each 8-item sequence. The forward model derived from Freesurfer’s fsaverage 3-layer mesh, and manually coregistered with the 32 scalp electrodes. The inverse model was fitted with a minimum norm estimate with MNE default parameters (lambda^2^=0.125, free dipole with normal component). The peak amplitudes and latencies (Fig. 4 B-C, E-F) were computed from the relative amplitude and relative latency of the maximal amplitude obtained for each source and each subject separately. The corresponding figures show these effects averaged across subjects.

### Statistics

Except if stated otherwise, all inferential statistical estimates derive from two-tailed second-level nonparametric analyses across subjects. Specifically, each decoding, encoding and source analysis was applied within each subject separately and led to a unique estimate of the effect size obtained across time samples (e.g. an R correlation coefficient for each subject). A second-level analysis was then applied across subjects to assess whether the distribution of these effect sizes was different from chance. This second-level analysis was either (1) a Wilcoxon test applied across subjects (in the case of a non-repeated analysis) or (2) a spatio-temporal cluster test applied across subjects (in the case of repeated measurements, such as decoding time courses or encoding spatio-temporal effects). The p-values of the decoding time-courses, TG matrix, sources estimates, and EEG topographies all refer to the p-value of the significant spatio-temporal cluster (Maris and Oostenveld, 2007) as implemented in MNE with default parameters (Gramfort et al., 2014). The error bars plotted in the figures correspond to the standard error of the mean across subjects.

### Modeling neural dynamics

We used a dynamical system framework to simulate the spatio-temporal dynamics evoked by one or several successive stimulations of fixed duration. A model example is illustrated in Fig. S4. Each model consists of a set of units *z* which each transforms the sum of their inputs *u* with a potentially nonlinear activation function (*f*):

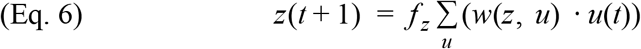

where *t* refers to time sample and *w(z, u)* refers to the connection from unit *u* to unit *z*. Except if stated otherwise, *f* is a linear activation function (i.e. *f(z) = z*). Each architecture consisted of a repeated pattern of connections between layers, each consisting of observable (*x*) and/or hidden (*y*) units connected through recurrent (*w*(*x*_*i*_, *x*_*i*_), *w*(*y*_*i*_, *y*_*i*_), *w*(*x*_*i*_, *y*_*i*_), *w*(*y*_*i*_, *x*_*i*_)), feedforward (*w*(*x*_*i*_, *x*_*i+1*_), *w*(*y*_*i*_, *y*_*i+1*_), *w*(*x*_*i*_, *y*_*i+1*_), *w*(*y*_*i*_, *x*_*i+1*_)) and feedback connections (*w*(*x*_*i*_, *x*_*i−1*_), *w*(*y*_*i*_, *y*_*i−1*_), *w*(*x*_*i*_, *y*_*i−1*_), *w*(*y*_*i*_, *x*_*i−1*_), Fig. 2C). In other words, each unit can only be connected to other units within the same layer (i), the layer above (i+1) or the layer below (i−1):

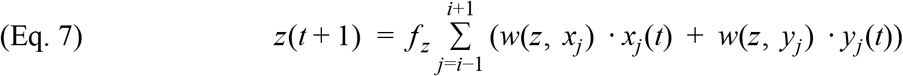

The simulated activity was analyzed with decoding and temporal generalization analyses after a linear random projection (F, normally distributed around 0, with s.d.=1) of the *x* units onto 32 noisy virtual sensors *X*:

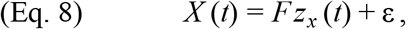

where X(t) is the activity of EEG channels at time t, z_x_ is the activity of observable units and ε is a gaussian noise (except if stated otherwise, s.d.=1).

This random projection and the subsequent decoding and TG analyses was repeated 15 times to mimic the analyses of 15 subjects.

Except in Figure 2A, units do not intend to simulate individual neurons, but aim to approximate the dynamical characteristics of the macroscopic electric field. In this view, some recurrent connections may be equally interpreted as an adaptation mechanism (i.e. activity reduction caused by a cellular mechanism) or as a lateral-inhibition mechanism (i.e. activity reduction caused by an inter-cellular mechanism). In either case, the hidden units *y* are designed to account for the possibility that some neural dynamics may be influenced by mechanisms that cannot be directly observed with EEG (e.g. adaptation does not generate an electric field, and the inconsistent orientations of the inter-neurons’ electric fields are not easily detected from distant EEG electrodes).

### Modeling representational structures

Following others (King and Dehaene, 2014; Kriegeskorte and Kievit, 2013), we equate neural representations and linearly-decodable information, on the basis that linear-readout can be performed by any neuron input with such information. In this view, the level of abstraction/complexity of a representation corresponds to the number of nonlinear computations applied to a given input. To investigate whether low-level representations are likely to remain linearly decodable from the highest processing stages of the visual system, we tested whether they would be linearly decodable from the layers of VGG-19, a deep convolutional neural network trained to label 5 million images with over 22,000 object categories. The training weights were downloaded from Keras (Chollet, 2015). VGG-19 was input with 1,000 Gabor patches similar to those used in our study. Thirty-two virtual EEG sensors were simulated by randomly projecting these high-dimensional activations from each layer separately (as in Eq. 8). Finally, ordinary-least-square regressions were fit on each activation layer of VGG-19, following the same procedure to the one used in our EEG decoding analyses (Fig. 5D, red). Finally, to verify that our approach dissociated low and high-level representations, we generated a series of arbitrary non-linear transformations of the stimulus angles (Fig. 4), and tested whether they were sequentially represented across the layers of VGG-19 and across the time samples of the EEG. Concretely, we investigated three hierarchical levels of representations defined as low=*f(angles)*, middle=*f(f(angles))* and high=*f(f(f(angles)))* respectively, where *f* is a normalized rectifier (*f*):

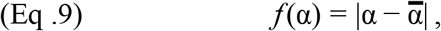

where *α* is the feature of a given sample *i*, and 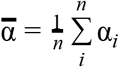 is the average feature across samples. For example, given uniformly distributed angles α ∈ [0, 2π] then, *f(0)* = *π*, *f(π)* = *0* and *f(2π)* = *π*.

## Acknowledgments

This work was supported by the European Union’s Horizon 2020 research & innovation program under the Marie Sklodowska-Curie Grant Agreement No. 660086 (J-R.K.), as well as by the Bettencourt-Schueller and the Philippe Foundations (J-R.K.). We are thankful to Gyorgy Buzsaki, Marisa Carrasco, Saskia Haegens, David Heeger, Lucia Melloni, David Poeppel and Eero Simoncelli and their teams for their valuable feedback. We are also grateful to the contributors of the MNE open-source package for their generous support.

## Author contributions

Conceptualization, J-R.K. and V.W.,; Experimental design: V.W. Methodology, J-R.K. and V.W; Software, J-R.K.; Data collection, V.W; Formal Analysis: J-R.K.; Resources, V.W. & J-R.K.; Data Curation, V.W.; Writing – Original Draft, J-R.K.; Writing – Review & Editing, J-R.K.; Visualization, J-R.K.; Project Administration, J-R.K.; Funding Acquisition, J-R.K.

