## supplementary for "The Human Brain encodes a Chronicle of Visual Events at each Instant of Time"

### Supplementary Materials

##### Inferring the beginning and end of significant clusters: what limitations?

In Fig. 1E, the cluster-corrected significance appeared to start at time 0. This early start is surprising given the necessary conduction delay between the retina and the cortex. This early statistical effect could thus be due to the imprecision of the cluster-corrected statistics. First, the corresponding decoding score is relatively close to chance at this time sample as compared to the rest of the cluster. Second, the spatio-temporal cluster test formally assesses whether there is a significant deviation from chance level, but is known to be limited in its ability to precisely identify when such effect starts (1).

##### Does activity reversal explain the sustained decoding scores?

Each decoder trained at a given time sample tends to generalize below chance level after the offset of the stimulus (Fig. 3A). These results suggest that the brain activity patterns coding for stimulus angles and deltas reversed between the onset and the offset response. Does this reversal suffice to explain the 1-second long decoding performance? To address this issue, we formalize two hypotheses:

H0: the stimulus elicits a brief series of responses, followed by pattern reversals.

H1: the stimulus elicits a long series of responses, accompanied by pattern reversals.

Both hypotheses predict above chance decoding performance after stimulus offset. However, H0 predicts that the decoders trained after stimulus offset would be equivalent to the decoders trained after stimulus onset (after a sign reversal). Conversely, H1 predicts that the decoders trained after stimulus offset would outperform the decoders trained after stimulus onset (after a sign reversal).

We thus compared diagonal decoders (i.e. trained and tested at the same time samples) to off-diagonal decoders (i.e. trained at time t and tested at time t+250ms, after sign reversal, Fig. S3A) . The results show that off-diagonal decoders were significantly above chance up to a ~1 s after the onset of the stimulus (blue thin line Fig. S3A), and thus confirm the significant pattern reversal. However, most diagonal decoders weakly but consistently outperformed off-diagonal decoders (Fig. S3. B-C).

These results suggest that the long-lasting decoding of stimulus angles and deltas are partly explained by pattern reversals, but are also contributed by additional propagating activity.

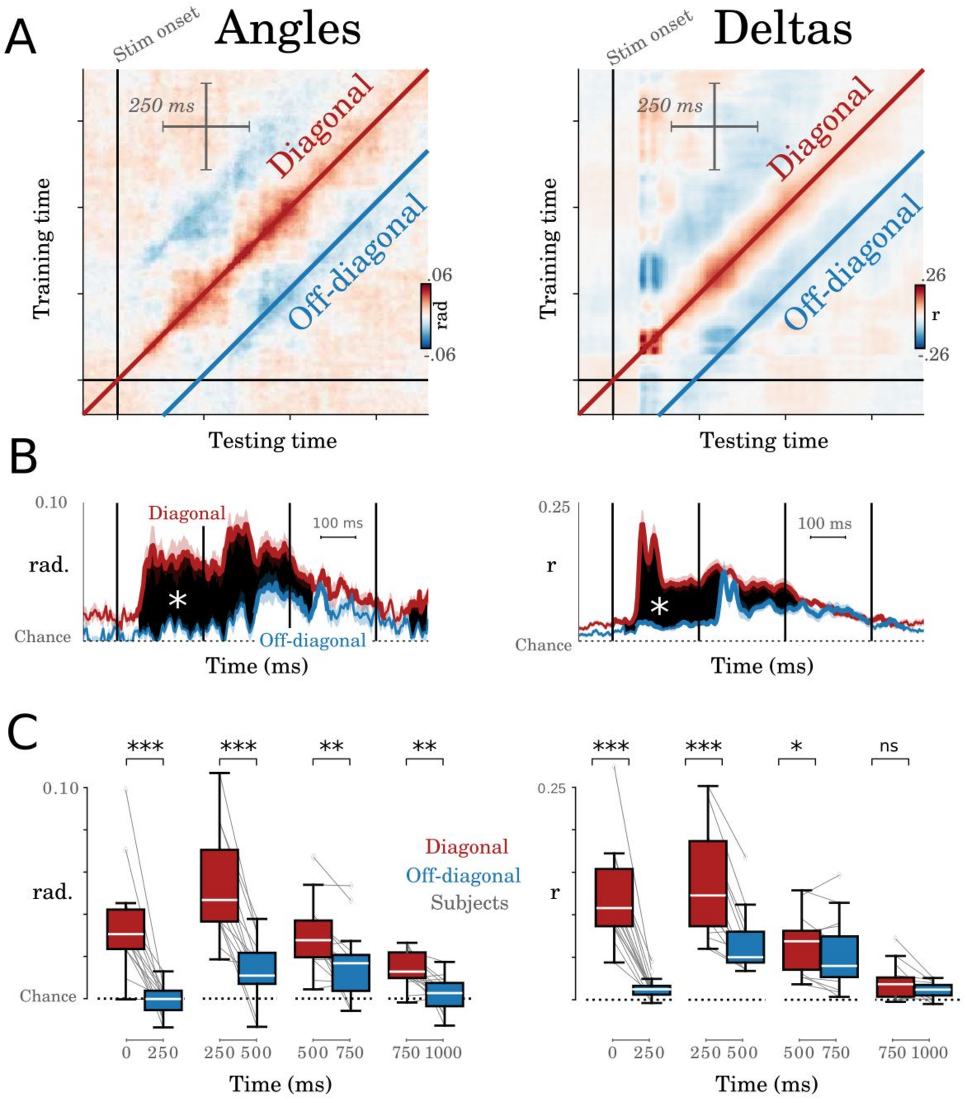


**Figure S1. A** Temporal generalization, identical to Fig. 3A. Diagonal decoding refers to the scores obtained from decoders trained and tested at the same time samples, and are used to isolate onset responses. Off-diagonal decoders refer to decoders trained at time t, and tested 250 ms later, in order to isolate offset responses. **B.** Diagonal and off-diagonal decoding scores (reversed in sign for clarity) as a function of time. Thick lines indicate when the decoders are statistically significant across subjects (cluster corrected). The shaded black region (with a star) indicates when the diagonal decoders significantly outperform off-diagonal decoders. **C.** Same data as B, after averaging decoding scores within four successive 250 ms windows. Stars indicate when diagonal decoders significantly outperform off-diagonal decoders (*: $p<0.05$; **:$p<0.01$; ***:$p<0.001$).

#### What biological mechanism underlie the dynamical system isolated in our study?

The below-chance decoding performance observed in Fig. 3. A-B, suggests that the EEG activity coding for the stimulus flipped in direction: e.g. the EEG channels that positively correlated with deltas at time *t*, negatively correlated with deltas at *t'*. Such activity reversal could be explained by a variety of biological mechanisms, such as (1) lateral-inhibition or (2) adaptation (Fig. 2-A). To investigate whether adaptation alone can also account for (1) the diagonal temporal generalization matrices and (2) the decodability of deltas, we simulated two models encoding the angle of the stimuli. The two models vary in the strength of their adaptation, i.e. in the amplitude of the inhibitory connection from *y* to *x* (Fig. S2).

(1) We first applied the same decoding and temporal generalization analyses as for our EEG recordings. As expected, the results show that the angle of the stimulus was decodable even after the stimulus onset (Fig. S2, black shaded lines). However, deltas could not be decoded from these simulations. In other words, a linear dynamical system that models sensory adaptation does not predict delta decoding.

(2) We then implemented temporal generalization (TG) analyses to investigate the dynamics of angle representations (Fig. S2. Bottom). As expected, the two TG matrices led to below-chance generalizations after stimulus offset. However, unlike our EEG analyses (Fig. 3A), none of the two matrices were characterized by a long diagonal pattern.

Overall, these results suggest that adaptation may partly explain some of our findings, such as the pattern reversal observed after stimulus offset, and the significant decoding of stimulus angles after stimulus offsets. However, the significant delta decoding and the long diagonal generalization scores empirically observed in our study suggest that additional neuronal mechanisms, identifiable with *in vivo* electrophysiology, may be at play.

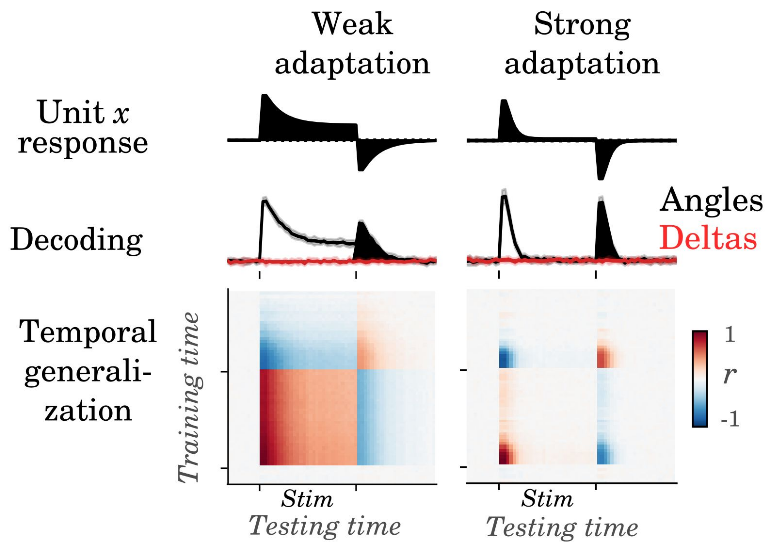


**Figure S2. Adaptation simulation.** Two models, with weak and strong adaptation respectively, were simulated (left versus right columns). **Top**. Unit $x$ responses illustrate the dynamics of an adaptive neuron in response to a stimulus (onset and offset marked by ticks). **Middle.** Decoding of stimulus angle$_n$ (black), and stimulus delta$_n$ (i.e. |angle$_n$ - angle$_{n-1}$|, in red) in response to a stimulus sequence.

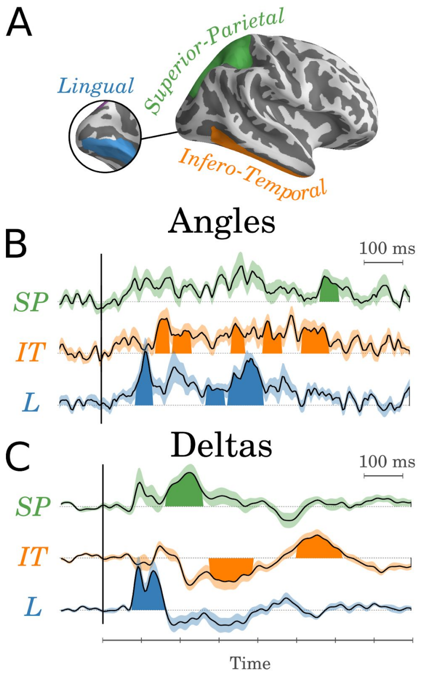


**Figure S3.** Average time-course observed in three anatomical regions of interest covering the ventral and dorsal visual pathways and derived from the Desikan-Killiany' cortical atlas (*2*) provided by Freesurfer (*3*): the bilateral lingual, inferotemporal and superior parietal cortices. **A.** Location of the three regions of interest. **B.** Average time course of angle representations in each region of interest (color coded) as revealed with encoding analyses. Shaded curves indicate significant clusters of activations across subjects. The dotted lines indicate the chance level. **C.** Analogous analysis to B for delta representations.

**Model search.**

Which architectures can account for our empirical observations? To address this question, we (1) implemented a grid-search analysis over architectures, connection weights and activation functions and (2) tested whether the resulting dynamics consisted of a similar spatio-temporal response.

(1) Our search considered hierarchical networks with one observable unit (*x*) and one hidden unit (*y*) per level *i*. Consequently, there are twelve possible connections: four recurrent recurrent ($w(x_i, x_i)$, $w(y_i, y_i)$, $w(x_i, y_i)$, $w(y_i, x_i)$), four feedforward ($w(x_i, x_{i+1})$, $w(y_i, y_{i+1})$, $w(x_i, y_{i+1})$, $w(y_i, x_{i+1})$) and four feedback possible connections ($w(x_i, x_{i-1})$, $w(y_i, y_{i-1})$, $w(x_i, y_{i-1})$, $w(y_i, x_{i-1})$, Fig. 2C). We presently define an "architecture" as the set of models whose feedforward, recurrent and feedback connections have identical signs. For example, two models A and B that each consists of a unique feedforward connection between unit $x_i$ and unit $x_{i+1}$ and whose weight is 0.75 and 1.00 respectively, belong to the same architecture: i.e. a positive feedforward architecture. However, a model C, that consists of a unique inhibitory feedforward connection between $x_i$ and $x_{i+1}$ (e.g. $w(x_i, x_{i+1})$=-0.50) belongs to a distinct architecture (i.e. negative feedforward architecture). In this view, there thus exists $3^{12}$ = 531,441 possible architectures. This set includes non-hierarchical networks (i.e. all feedforward connections are set to 0) and fully observable networks (i.e. there are no connection from *y* units). For each connection, we tested 21 possible values, linearly distributed between -1 and 1. Finally, to extend our model search to nonlinear dynamics, we also search across four monotonic activation functions:

(Eq. 10) Linear: $f(z) = z$

(Eq. 11) Relu: $f(z) = max(z, 0)$

(Eq. 12) SatRelu: $f(z) = min(max(z, 0), 1)$

(Eq. 13) SatLin: $f(z) = min(max(z, -1), 1)$

The above activation functions were applied independently for *x* and *y* units, and thus lead to $4^2$ =16 combinations of activation functions per architecture (e.g. *Linear(x)+Linear(y)*, *Linear(x)+Relu(y)* … *Satlin(x)+Satlin(y)*).

Overall, the total search thus spans over $4^2 \times 21^{12} > 10^{17}$ distinct models. All models were implemented with 10 hierarchical levels, and simulated over 120 time samples, with a constant input between time 30 and 60. The input was connected to the network following the feedforward weights. Because we are interested in finding the simplest architectures that can account for our empirical result, we only report the successful architectures with the lowest number of non-zero connections.

(2) Each model was assessed on its ability to account for the three main findings identified in our EEG study: (2.1) Stimulus onset evokes a transient traveling wave ('onset'), (2.2) Stimulus offset evokes a transient traveling wave with opposite amplitude ('offset') and

(2.3) the phase (i.e. 'width') of these waves increases across levels ('increasing maintenance'). We quantified these properties for the observable units *x*. The *y* units being hidden, they could thus have any dynamical response.

(2.1) **Onset.** A unit was considered to be marked by an 'onset' if its maximum value $M$ reached at time $t_M$ was positive, if $t_M$ was before stimulus offset, and if all values between stimulus onset and $t_M$ increased over time:

(Eq. 14) $$(x(t_M) > 0) \cup (t_M < offset) \cup (\dot{x}(t) \geq 0,\ t \in [onset,\ M])$$

A network was considered to generate an 'onset' traveling wave if all of the x units were marked by an onset at each level.

(2.2) **Offset.** A unit was considered to be marked by an offset if its minimum value $m$ reached at time $t_m$ was negative, and if all values between $m$ and the end of the simulation increased towards 0:

(Eq. 15) $$(x(t_m) < 0) \cup (t_m > offset) \cup (\dot{x}(t) \geq 0,\ t \in [m,\ end])$$

A network was considered to generate an 'offset' traveling wave if all of the $x$ units were marked by an offset at each level.

(2.3) **Increasing maintenance.** A maintenance 'half-life' was estimated for each level by estimating by the delay $t_h$ it takes a unit to reach half of its maximum value $M$: $x(t_h) = x(t_M)/2$ . Half-lives were estimated if all values between $t_M$ and $t_m$ decreased towards 0:

(Eq. 16) $$(x(t) \geq 0,\ t \in [M,\ offset]) \cup (\dot{x}(t) < 0,\ t \in [M,\ m])$$

The network was considered to have increasing maintenance if half-lives increased across levels:

(Eq. 17) $$\frac{dh}{di} \geq 0$$

Out of the 531,441 architectures, two architectures matched our empirical findings with no more than four connections/parameters. These two architectures could match our EEG recordings with either linear and/or non-linear activation functions. In other words, non-linearities did not appear necessary to explain our empirical observations.

#### Could subjects perform the task without maintaining the visual stimuli?

As detailed in the methods, the subjects' task consisted in determining whether the orientation of the stimuli fell, on average, closer to the cardinal axes or closer to the diagonal axes. A response had to be provided at the end of each 8-item sequence.

To demonstrate that this task can be performed without maintaining the orientation of each stimulus, we simulated a two-layer neural network (Fig. S4A). The first layer of this model is referred to as "sensory layer". Its connectivity to the input stimuli is designed such that it encodes the orientation of each stimulus. Specifically, it connects a one-hot-encoder of the orientations $\alpha$ (i.e. one input dedicated to each orientation) to each sensory neuron tuned to orientation $\mu$ of the sensory layer following a von Mises function $g$:

$$(\text{Eq. 18}) \qquad g(\alpha, \mu) = \kappa\, e^{\kappa \cos(\alpha - \mu)} \times \frac{2\pi}{i_0(\kappa)}$$

where κ (which controls the encoding precision) is fixed to 2, such that $g(0, 0) = 1$ and $g(\pi, 0) = 0$, and where $i_0$ is the modified Bessel function provided by the Scipy package (Fig. S4B, orange line).

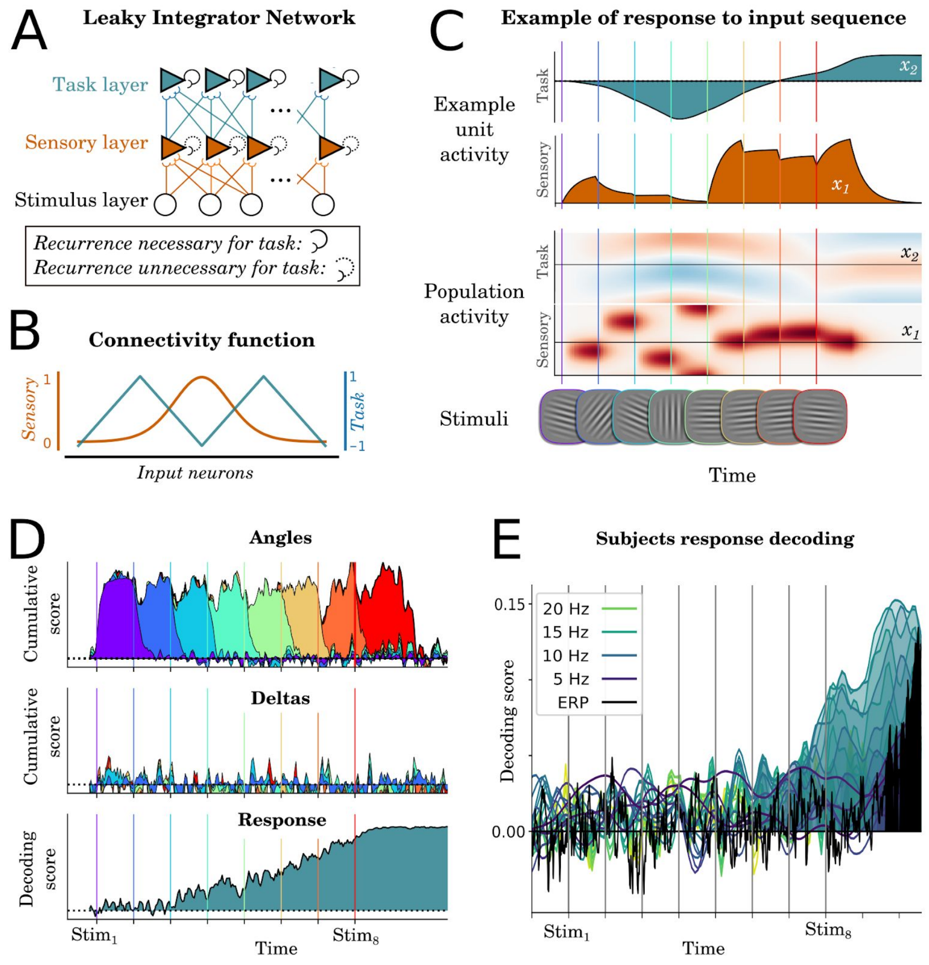


**Figure S4. A.** A two-layer neural network with fully observable units can perform the task without encoding a chronicle of visual events at each time instant. The sensory neurons (orange) encode the angle of the stimulus with or without a recurrent connection whereas the task neurons transform and accumulate these sensory representations to perform the task. **B.** The connectivity functions used to encode (i) the sensory representations from a one-hot-encoder of the angles (von Mises function, in orange) and (ii) the task representations from the sensory representations (sawtooth function, in gray) **C.** Simulated neuronal responses to a random sequence of 8 stimuli. The shaded lines illustrate the activity of (i) a sensory neuron encoding horizontal orientations ($x_1$) and of (ii) a task neuron encoding the cardinal axis ($x_2$). The heat map below illustrates the entire population response to the eight stimuli (red=positive response, white=no response, blue=negative response). Vertical bars indicate stimulus onsets **D.** Cumulative decoding of the simulated neuronal activity projected onto 32 virtual sensors as a function of time (x-axis) and stimuli (color-coded) for both angles, deltas, and trial response. **E.** EEG decoding performance of subjects' responses as a function of time (x-axis) and frequency (black: raw evoked potential, dark blue: EEG power at 5Hz, light green: EEG power at 20 Hz). Shaded lines indicate significant clusters of decoding scores across subjects.

The second layer of this model is referred to as "task layer". Its connectivity to the sensory layer is designed such that this layer encodes and continuously updates the response subjects were supposed to give in a given trial. Specifically, the task layer encodes whether the average orientation in a given trial falls closer to the cardinal axes or to the diagonal axes. Its connectivity from the sensory layer is defined by a sawtooth function with two cycles over the sensory neurons (Fig. S4B, blue line). Critically, strong recurrent connections (here $w(x_i, x_i)=0.999$) is necessary for this layer to accumulate task-related information.

Finally, we randomly projected all of these simulated neurons onto virtual noisy EEG sensors and applied the same decoding analyses to those employed for the empirical recordings (see methods). The results, summarized in Fig. S4D confirm that (1) the sensory information related to successive stimuli need not be maintained over time, (2) the deltas between successive stimuli are not decodable in this simple model and that (3) task-related information can nonetheless be decoded over the course of an 8-item sequence.

Finally, to investigate the plausibility of the task-layer, we applied a linear decoder at each time sample time-locked to the beginning of each 8-item sequence, in order to determine whether task-related information followed the prediction of our model. The linear classifier (Logistic regression, as provided by the Scikit-learn package) was fit with either the raw evoked potential (Fig. S4-E, black) or the instantaneous power in a given frequency band (Fig. S4E blue and green lines), as estimated with a Morlet wavelet decomposition (5 cycles, frequency ranging from 3 to 24 Hz, separately for every 1 Hz). The results confirmed that task-related information could be increasingly decoded from the end of the fifth stimulus and until the end of the trials.

Together, the temporal profile of our simulations and of the empirical task decoding scores suggest that the task is performed by a set of neurons distinct from those encoding and maintaining successive visual events. Overall, these results suggest that our ability to simultaneously decode successive visual stimuli at each time instant is not trivially explained by subjects' task.

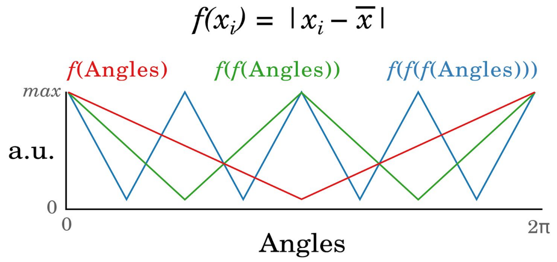


**Figure S5.** Illustration of increasingly complex features generated by the *f* function.

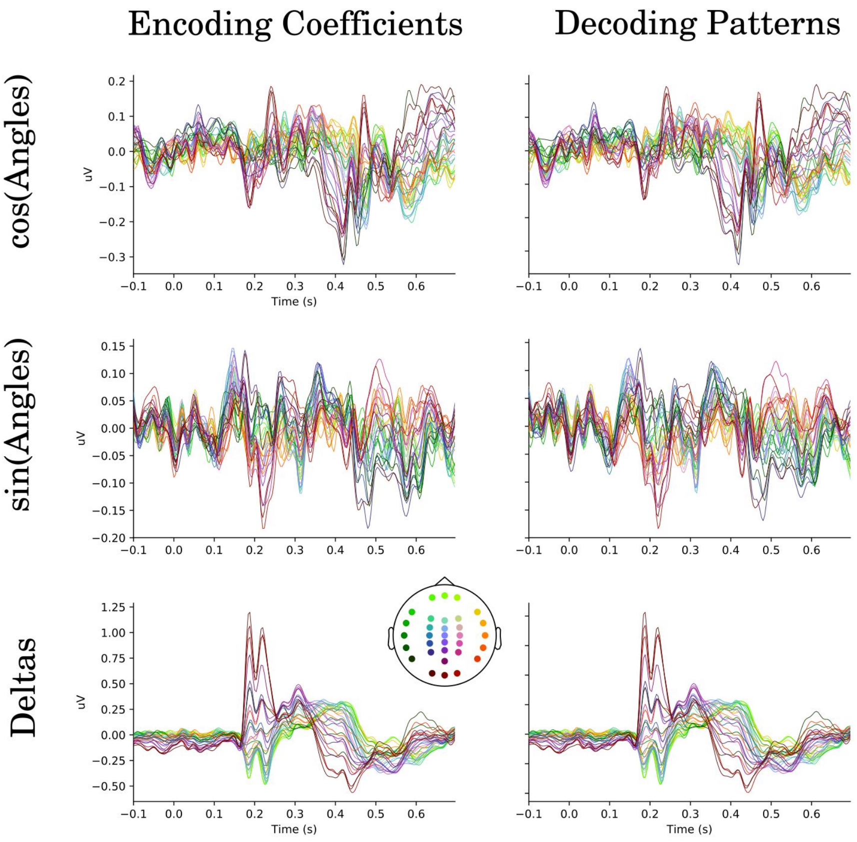


**Figure S6.** Encoding coefficients and decoding patterns averaged across subjects. Each line indicates an EEG channel (color-coded by its spatial position).

#### Supplementary methods: simulation of biological mechanisms.

Adaptive exponential neurons were modeled following Brett and Gestner 2005. Specifically, their activity was modeled as a two-dimensional nonlinear dynamical system:

(Eq. 19) $$\dot{V}_m = (g_L(E_L - V_m) + g_L\Delta_t \exp((V_m - V_T)/\Delta_t) - w + I) \times \frac{1}{C}$$

(Eq. 20) $$\dot{w} = a \times (V_m - E_L) - w \times \frac{1}{\tau_w}$$

where $V_m$ measures the membrane potential, $w$ modulates adaptation, and C (281 pF), $g_L$ (30 nS), $E_L$ (-70.6 mV), $V_T$ (-50.4 mV), $\Delta_t$ (2 mV), a (4 nS), and $\tau_w$ (144 ms) are fixed constants derived from Brett and Gestner 2005.

A spike was generated when the membrane potential was superior to $V_{cut}$, where

(Eq. 21) $$V_{cut} = V_T + 5 \times \Delta_t$$

After each spike, the membrane potential was reset to -70.6 mV, and $w$ was incremented by 0.0805 nA.

Ten thousand neurons were simulated for 1 s. Each neuron received a Poisson spiking input, where each input spike increased the membrane potential by 10 mV to mimic random input from distant neurons. The 'stimulus activity' lasted for 233 ms and corresponded to a 0.9 nA convolved with a von-Mises distribution (as in Fig. S4B) that mimics the tuning of a given orientation.

Excitatory-inhibitory dynamics were approximated as mean-field activity potentials generated by a Cowan model. Specifically, a population of excitatory and inhibitory neurons (A) whose saturating activity is controlled by a sigmoid (σ) was connected with one another through a connectivity matrix $W$.

(Eq. 22) $$A(t+1) = \sigma(W \cdot A(t) + I(t))$$

Recurrent excitatory connections, Recurrent excitatory connections, and excitatory to inhibitory connections were set to 1, 1, and -1 respectively.

EEG is not sensitive all neuronal activity, but to the post-synaptic potentials input to the vertically-aligned dendrites of excitatory neurons. Consequently, in Figure 2A the x-unit corresponds to the average inputs to the excitatory neurons.

The input amplitudes to excitatory and inhibitory units were manually set to 0.15 and 0.10 in order to generate a dynamics similar to the spiking rate of adaptive exponential neurons.
